# Barcoding genetically distinct *Plasmodium falciparum* strains for comparative assessment of fitness and antimalarial drug resistance

**DOI:** 10.1101/2022.04.05.487250

**Authors:** Manuela Carrasquilla, Ndey Fatou Drammeh, Mukul Rawat, Theo Sanderson, Zenon Zenonos, Julian C Rayner, Marcus CS Lee

**Affiliations:** Wellcome Sanger Institute, Wellcome Genome Campus, Hinxton, UK; Max Planck Institute for Infection Biology, Berlin, Germany; Medical Research Council Unit The Gambia, at the London School of Hygiene and Tropical Medicine, Banjul, The Gambia; The Francis Crick Institute, London, UK; Biologics Engineering, Early Oncology, AstraZeneca, Granta Park, Cambridge, UK; Cambridge Institute for Medical Research, University of Cambridge, Cambridge, UK

## Abstract

The repeated emergence of antimalarial drug resistance in *Plasmodium falciparum*, including to the current frontline antimalarial artemisinin, is a perennial problem for malaria control. Nextgeneration sequencing has greatly accelerated the identification of polymorphisms in resistance-associated genes, but has also highlighted the need for more sensitive and accurate laboratory tools to profile current and future antimalarials, and to quantify the impact of drug resistance acquisition on parasite fitness. The interplay of fitness and drug response is of fundamental importance in understanding why particular genetic backgrounds are better at driving the evolution of drug resistance in natural populations, but the impact of parasite fitness landscapes on the epidemiology of drug resistance has typically been laborious to accurately quantify in the lab, with assays being limited in accuracy and throughput. Here we present a scalable method to profile fitness and drug response of genetically distinct *P. falciparum* strains with well-described sensitivities to several antimalarials. We leverage CRISPR/Cas9 genome-editing and barcode sequencing to track unique barcodes integrated into a non-essential gene (*pfrh3*). We validate this approach in multiplex competitive growth assays of three strains with distinct geographical origins. Furthermore, we demonstrate that this method can be a powerful approach for tracking artemisinin response as it can identify an artemisinin resistant strain within a mix of multiple parasite lines, suggesting an approach for scaling the laborious ring-stage survival assay (RSA) across libraries of barcoded parasite lines. Overall, we present a novel high-throughput method for multiplexed competitive growth assays to evaluate parasite fitness and drug response

## Introduction

Antimalarial drug resistance repeatedly emerges first in certain regions of the world, a phenomenon that has been linked to particular genetic backgrounds that better tolerate resistance-associated polymorphisms^1–4^. The independent origin of chloroquine and pyrimethamine resistance provided an example, in which beneficial mutations to drug pressure were rapidly fixed in Southeast Asia (SEA) and South America (SA), before subsequent dissemination to Africa^5–8^. Even though these resistance mutations could have emerged locally in Africa, where both parasite diversity and burden is higher, a high fitness cost likely prevented them from being maintained: a rapid decrease in frequency of resistance alleles in Malawi as a consequence of chloroquine withdrawal supported the hypothesis that evolutionary forces were acting differently on these populations than in SEA and SA^9^. Accurately and sensitively studying the impact of resistance-associated polymorphisms on both resistance and fitness is therefore key to defining the fitness landscapes of parasite populations globally.

The impact of drug resistance on parasite fitness has proven difficult to quantify in a laboratory setting. A major constraint on throughput is the fact that most fitness comparisons involve performing head-to-head competition assays with just two strains, a test line and a reference parasite, for which multiple readouts have been employed. One approach to detect the relative proportion of two alleles in a mixture is pyrosequencing, which uses the pyrophosphate released by polymerase incorporation of specific nucleotides to drive a luciferase-based enzymatic reaction. This light emission allows accurate quantification of the incorporation of nucleotides by using allele-specific primers. By this approach, the contribution of drug-resistant alleles can be determined by co-culturing resistant and sensitive clones^2,10^. Another approach to head-to-head competitive assays is to use a non-fluorescent line of interest co-cultured with a GFP-competitor line, with flow cytometry reporting changes in relative proportion^2,11,12^. Lastly, quantitative PCR has also been employed to measure relative abundance of co-cultured *P. falciparum* lines, by using probes detecting different alleles of drug resistance genes^12^. These methods have highlighted the importance of measuring fitness in understanding the complexity of drug resistance in *P. falciparum* parasites, however none allows for simultaneous measurement of large numbers of lines.

The ability to track the growth of many cell lines in parallel has been an important tool for phenotyping in model organisms such as yeast^13^, but is relatively new to apicomplexan parasites. Recent examples of this are the application of barcode sequencing (BarSeq) for high throughput growth phenotyping in *P. berghei*, using systematic reverse genetic approaches to generate knockout mutants with barcodes for fitness tracking^14^. Another recent study by Sidik *et al*. (2016) used CRISPR/Cas9-based genome editing to target all genes across the genome of *Toxoplasma gondii*, with the unique gRNA for each gene acting as a barcode^15^. Together, these studies measured the fitness of particular knockout mutants by tracking unique short DNA tags by next-generation sequencing (NGS), however the aim was to understand gene essentiality, rather than intrinsic parasite fitness, or drug response. Scalable approaches to understand the fitness impact of specific variants, either in the context of *in vitro* drug selections, natural variants in field isolates, or the influence of genetic background on buffering resistance driver mutations will be important as next generation antimalarials are developed and deployed.

Here, we develop a barcode tagging approach for *P. falciparum*, which lacks the high transfection efficiencies of *P. berghei* or *T. gondii*, and apply it specifically to the study of drug resistance and parasite fitness. We used CRISPR-editing to insert short barcode cassettes at a nonessential safe-harbour locus, the pseudogene *Pfrh3*, resulting in stable maintenance and segregation of a single-copy tag for each line. Critically, all barcodes were inserted at the same genomic site, and flanked by the same sequences, meaning that multiple tagged lines could be pooled, grown together under different selective conditions, and their relative proportions quantified using a single PCR reaction followed by next-generation sequencing. As proof-of-principle, we barcoded genetically and phenotypically diverse parasite strains from different geographic regions and with different drug resistance profiles. The barcoded lines were pooled to perform competitive assays to measure both inherent growth differences as well as drug response to antimalarial compounds. We also performed ring-stage survival assays (RSA) for artemisinin resistance using barcode sequencing as a readout, demonstrating the potential for complex assays to be carried out on laboratory strains, or the progeny of genetic crosses, using pooled high-throughput screening.

## Results

### Generating a panel of uniquely barcoded parasite clones

To develop a scalable approach for stable barcoding of different parasite lines, we generated a library of 94 different barcoded donor vectors encoding an 11 bp barcode sequence within a 120 bp cassette. This barcode library was assembled into a pCC1 vector with flanking ~1kb homology regions of the pseudogene *Pfrh3* (PF3D7_1252400)^16^, resulting in a pool of up to 94 different donors, as recently described^17^. To generate stable integrated barcodes, we generated an *rh3*-targeting Cas9/sgRNA expression plasmid, replacing the existing hDHFR cassette with the yeast *yDHODH* selectable marker to allow for co-selection of Cas9-gRNA and donor plasmids (Fig. 1A)^18,19^. The pool of barcoded donors was cotransfected with a Cas9/gRNA plasmid into three genetically diverse strains of *P. falciparum* - the reference 3D7 parasite, multidrug-resistant V1/S from Vietnam, and a recent Cambodian isolate (PH0212-C) harbouring the C580Y mutation in *Pfkelch13* associated with artemisinin tolerance (Fig. 1D)^20^. Primers targeting the flanking regions of the barcode insertion site within *rh3* were used to generate amplicons for barcode sequencing (BarSeq) using next-generation sequencing.

**Figure 1.**
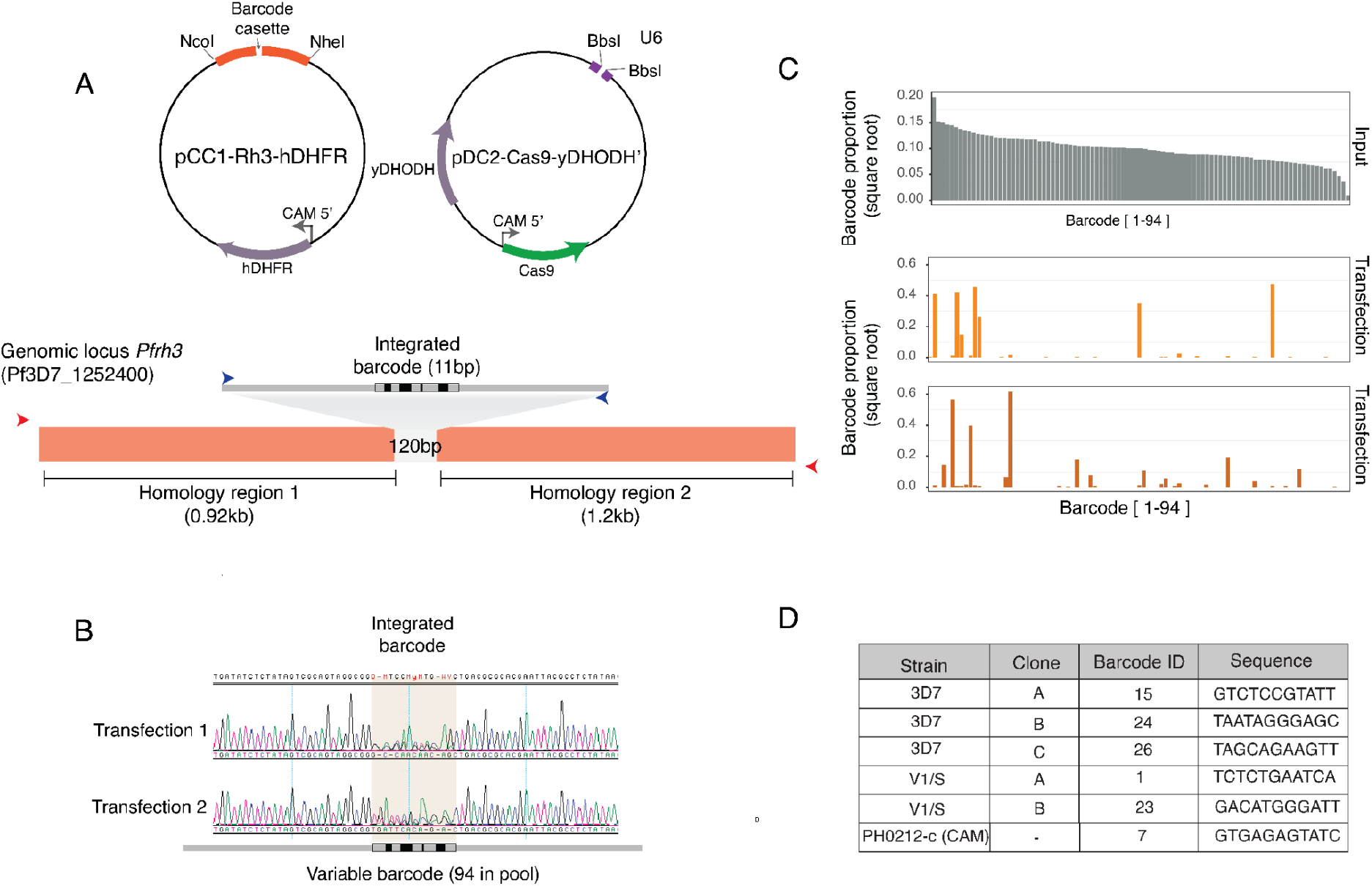
(A) Graphical representation of pCC1 donor plasmid and pDC2 Cas9 plasmid co-transfected for integration of a barcode cassette flanked by homology regions of *pfrh3* for homologous recombination. Red arrows in the bottom panel indicate the first PCR product generated from outside the homology regions to avoid episomal amplification. Blue arrows correspond to the second PCR inside the barcode cassette used for generating Illumina-compatible libraries. (B) Sanger sequencing chromatogram showing integrated barcodes after co-transfection of a pool of barcoded donor plasmids together with the pDC2-Cas9 vector. (C) Next Generation Sequencing of the integrated barcode region showing the input vector pool (top panel, grey) and the integrated barcodes, represented by the orange bars, for two independent transfections of 3D7. (D) Barcoded clones from the three strains used, the clone ID, the barcode ID from the pool of 94 barcodes, and the corresponding barcode sequence.

We first wanted to evaluate the complexity of barcode integration in a bulk culture of 3D7. Two independent transfections in this strain were initially assessed by Sanger sequencing, which revealed diverse nucleotide compositions at the barcode integration site within the bulk culture, consistent with multiple editing events having successfully taken place (Fig. 1B). BarSeq analysis of these transfections showed that 7-9 unique barcodes were recovered, confirming multiple different barcoded lines were present in the bulk population of transfectants (Fig. 1C). There was no strong bias towards barcodes that were more highly abundant in the original vector input pool (Fig. 1C, upper panel). Limiting dilution cloning was carried out to recover unique 3D7 barcoded lines. We then performed the same pooled transfections into strains CAM (Cambodia) and V1/S, observing a much lower complexity of the bulk population compared to 3D7, with only 1 and 2 barcodes recovered after cloning, respectively. Collectively, we selected six uniquely barcoded clones in total from these different genetic backgrounds (Fig. 1D) for use in downstream co-culture experiments.

### Change in barcode proportion reveals growth and drug sensitivity phenotypes

To experimentally test the pooled competition approach, we mixed all six lines in equal proportion, in three independent pools (Fig. 2A) to compare their intrinsic growth phenotypes. Importantly, these competition assays included more than one uniquely barcoded clone for both the 3D7 and V1/S strains. These act as replicates for each other, allowing for internal biological controls to be compared simultaneously for these two strains. The independent pools were cultured for 18 days in the absence of antimalarials, with samples of genomic DNA being extracted every 2-3 days for BarSeq (Fig. 2B). To generate growth curves derived from absolute read counts, we measured relative abundance of the different strains as the proportion of each barcode at a given time point. Overall, in the absence of drug pressure, the wild type 3D7 lines outcompeted the multidrug resistant V1/S lines but not the CAM parasite (Fig. 2B and 3A). Notably, the replicates for the 3D7 and V1/S lines within a pool were in each case internally consistent and showed no appreciable difference in growth between them, as expected (Fig. 2B).

**Figure 2.**
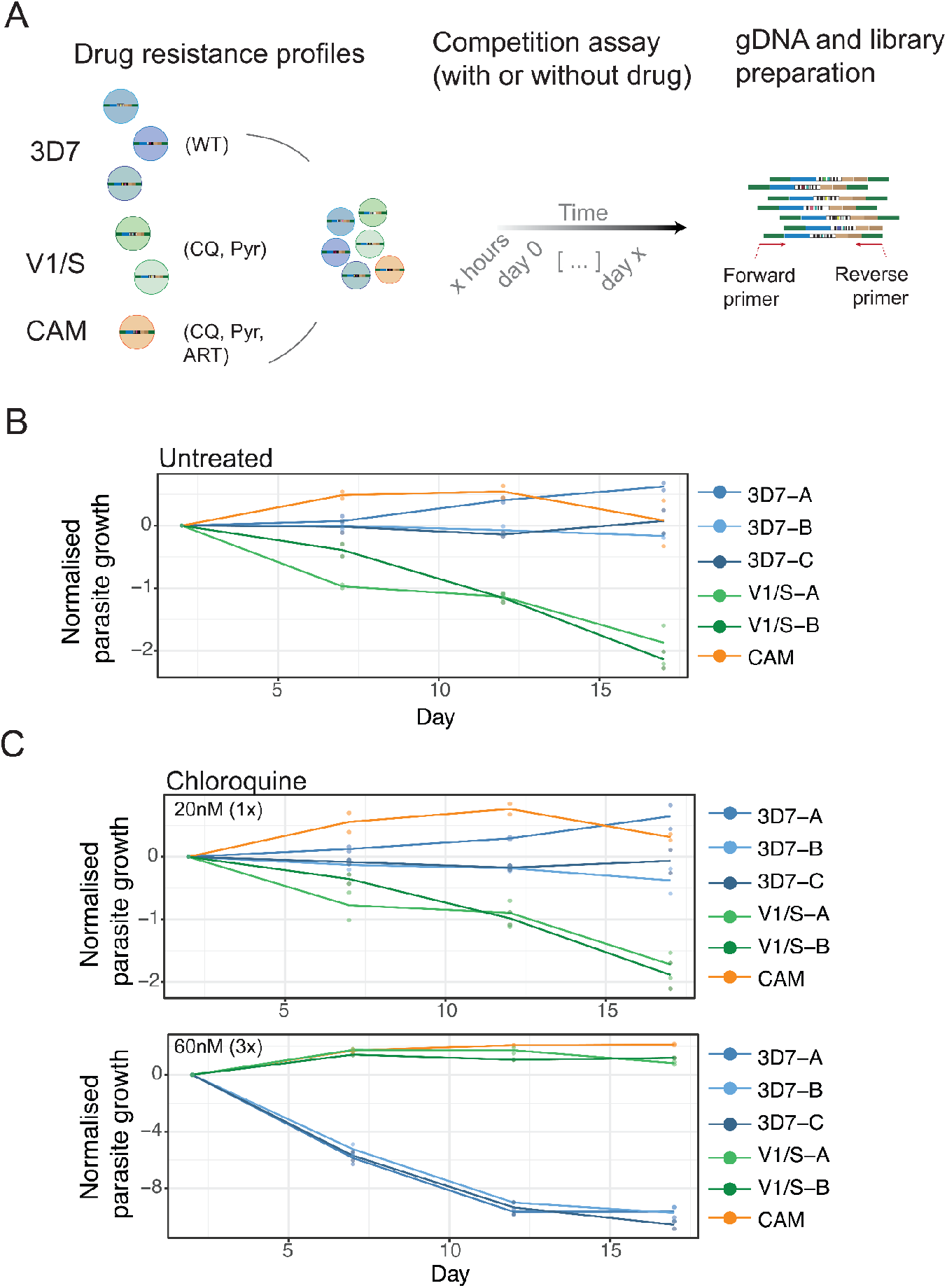
Competition assays to assess fitness and drug response. (A) General experimental design for BarSeq. Genetically diverse strains with different drug sensitivity profiles (CQ-chloroquine, Pyr-pyrimethamine, ART-artemisinin) are pooled and then grown together for multiple cycles in the presence or absence of drug selection. Samples are taken at multiple timepoints, genomic DNA extracted, and nested PCR reactions carried out to amplify the barcodes integrated into the *Pfrh3* locus of each strain. The relative proportion of each barcode is then quantified using NGS. (B-C) Competition assays lasting 18 days with two biological replicates (ie. two independent pools of all six strains, grown in parallel). Following BarSeq, the relative proportion of each strain was normalised to its abundance at the first time point and log2-transformed. (B) shows growth of the pools in the absence of selection, while (C) shows growth in the presence of two different chloroquine concentrations, corresponding to approximately 1 × or 3× the IC_50_ of the sensitive strain 3D7.

**Figure 3.**
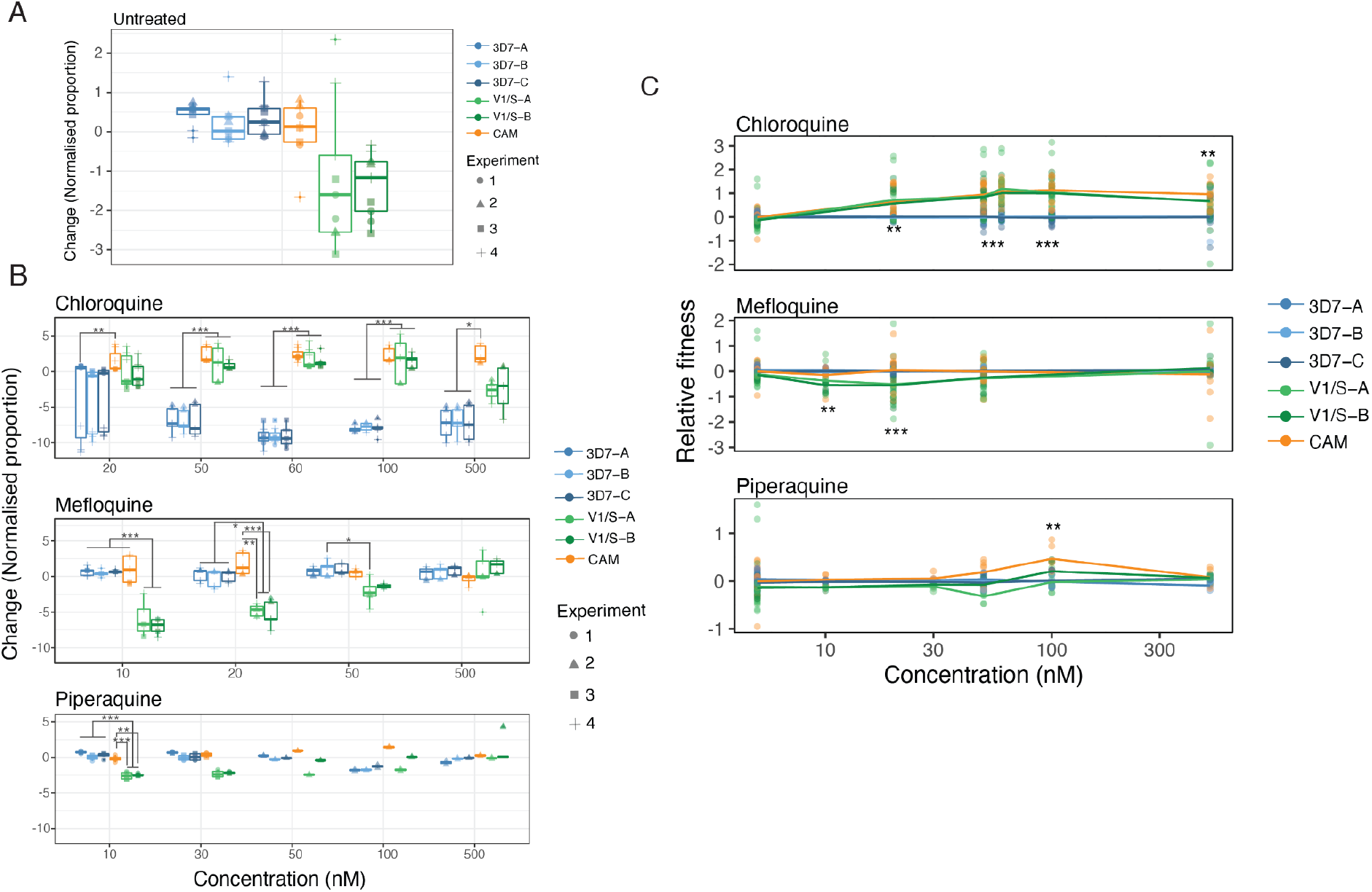
Drug response and relative fitness of barcoded strains. (A-B) Competition assay of barcoded strains in four independent experiments (9 biological replicates). Normalised proportions between the end time point and the first time point were log2-transformed (y-axis) and significant differences were calculated with a one-way ANOVA and TukeyHSD test for multiple comparisons of means. Competition assay in the absence of drugs showed consistent growth disadvantage of two barcoded clones of V1/S in the absence of any drug selection (untreated) (A). (B) Change in normalised proportion in barcoded strains to determine their antimalarial response under three drugs: chloroquine (top), mefloquine (middle) and piperaquine (bottom) and at increasing concentrations starting from the lowest IC_50_ of the three strains (Supplementary Table 1) and ramping up. (C) Change in fitness relative to the control strain 3D7 including all biological replicates. The x-axis represents the increasing concentration of chloroquine (top), mefloquine (middle) and piperaquine (bottom), normalised to the growth rate of the untreated control for the three independently barcoded lines 3D7 (3D7-A, 3D7-B and 3D7-C). Significant differences relative to 3D7 were determined at each timepoint using ANOVA and TukeyHSD (starred).

We next asked whether BarSeq could capture changes in growth rates in the presence of antimalarial drugs, detecting changes in sensitivity at increasing drug pressure. Drug concentrations were based on the IC_50_ of the 3D7 sensitive strain (Fig. 2A and Supplementary Table 1). Both the V1/S and CAM strains have the chloroquine-resistant allele of *pfcrt*, and as chloroquine pressure was increased from 1×IC_50_ to 3×IC_50_ of the sensitive strain, the growth advantage of 3D7 was reversed and both CAM and V1/S rapidly dominated the parasite population (Fig. 2C).

We further expanded our analysis to other clinically used antimalarials, mefloquine and piperaquine. Mefloquine has been used in combination with artesunate since significant resistance in Thailand and Cambodia emerged following its use as a monotherapy^21,22^. However, recent and rapid propagation of resistance to both drugs has rendered the use of this ACT completely ineffective in some areas of Southeast Asia^23,24^.

When the pooled strains were selected using mefloquine at 10nM, a concentration below the IC_50_ for 3D7 (32nM), we observed a significant hypersensitivity phenotype for V1/S (Fig. 3B and Supplementary Fig. 1A), consistent with our experimental values obtained from standard 72h dose-response assays (Supplementary Table 1) as well as previous findings (Duraisingh et al. 2000). Thus, mefloquine exposure inverts the 3D7-V1/S fitness relationship observed under chloroquine pressure (Supplementary Fig. 1A). At low levels of piperaquine (10nM-30nM), no clear change in overall profile was observed compared with untreated (Supplementary Fig. 1A), with V1/S being gradually outcompeted, presumably due to its inherently lower growth rate (Figure 2A) rather than drug response. Higher piperaquine concentrations appeared to provide a modest but not significant growth advantage for the CAM line, and the inherent growth disadvantage of V1/S was also partially negated under piperaquine pressure (Fig. 3B and Supplementary Fig. 1B). These results are consistent with the two-fold higher IC_50_ for piperaquine of both the CAM and V1/S lines relative to 3D7 (Supplementary Table 1), and support an overall advantage of the CAM strain at higher piperaquine concentrations attributable to higher fitness relative to V1/S and higher IC_50_ relative to 3D7. Treatment at the highest concentrations of mefloquine and piperaquine (500nM) resulted in a collapse of the growth for all lines (Fig. 3B and Supplementary Fig. 1B) as this concentration was well in excess of 10×IC_50_ for all lines. In contrast, 500nM chloroquine was less than 2×IC_50_ of V1/S and slightly below the IC_50_ for CAM, and thus still yielded meaningful data (Supplementary Fig. 1B). We then calculated the relative growth per day for each barcode at each timepoint, normalising each data point to the mean value from the three individually barcoded clones of 3D7. By calculating the mean to all the individual barcoded replicates measured (Fig. 3), we were able to establish fitness relationships between strains. These relationships were more easily visualised by plotting the change in fitness relative to 3D7 over different drug concentrations, and allowed us to statistically compare strains at a given concentration (Fig. 3C). Overall, these results illustrate the capacity of the BarSeq assay to capture how relative fitness is modulated by drug sensitivity, and the potential of the assay to test multiple drugs and drug concentrations in parallel.

### Artemisinin response measured by barcode sequencing

There is a pressing need to more fully understand how parasite populations, particularly in Southeast Asia, are adapting to the frontline antimalarial artemisinin. In addition to distinct mutations in the *pfkelch13* gene, of which the most common is C580Y, there is also evidence that the genetic background is instrumental in buffering potential fitness costs of different *pfkelch13* alleles^2,3^. Efforts to dissect the contribution of the genetic background in compensating for the acquisition of artemisinin-resistance are now being aided by the use of genome editing to insert *pfkelch13* alleles into distinct strains, and genetic crosses using the humanised mouse model^25,26^. However a major bottleneck is the subsequent large-scale phenotyping of the resulting lines, both in terms of fitness cost incurred in the absence of drug and individual artemisinin tolerance, which is conventionally measured using the laborious ring-stage assay (RSA) method^27^. We took advantage of our pool containing a Cambodian isolate with the PfKelch13-C580Y allele to explore whether the multiplex barcode approach could be exploited to phenotype artemisinin sensitivity.

Continuous exposure of the parasite pool to 10nM dihydroartemisinin (DHA), the active metabolite form of artemisinin, using similar conditions to the experiments above did not yield significant differences between strains (Fig. 4A). Artemisinin has a short half life, which combined with the frequency of treatment failure^28^, has resulted in the need to develop assays to accurately measure treatment failure due to resistance. Standard inhibitory assays have failed to do so, as they do not show a significant shift in the resistant lines that is well correlated with the slow-clearance phenotype in clinical cases. As an alternative, the Ring Stage Survival Assay (RSA) involves treatment of early rings, a stage shown to be resilient in slow-clearing clinical isolates, at a high concentration of DHA^27^. We first tested a simplified and less laborious approach, performing daily 3h pulses of unsynchronised cultures with 10nM DHA over a period of 17 days. However, no enrichment of the Kelch13-mutant strain was observed (Supplementary Fig. 2A, B).

**Figure 4.**
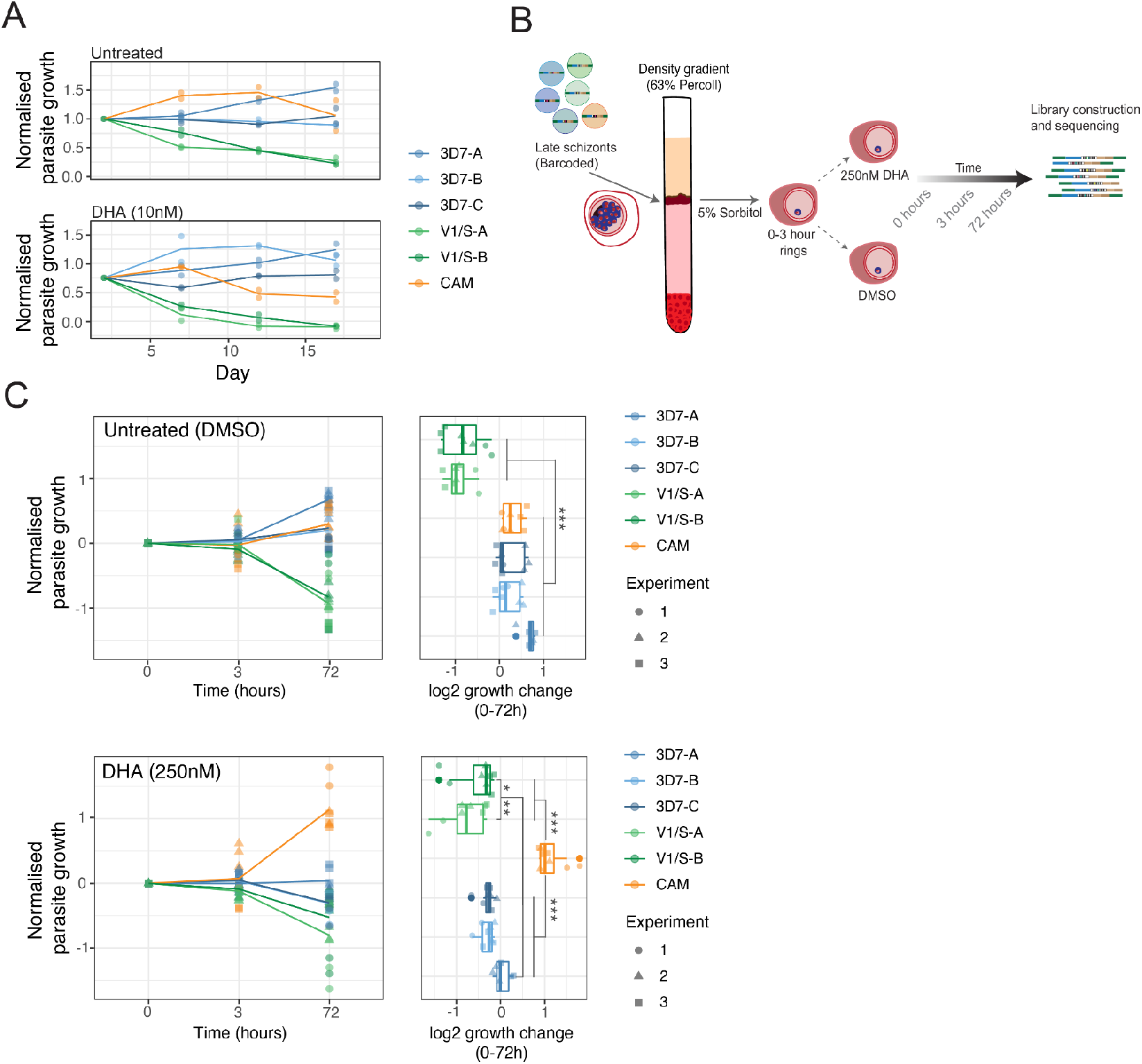
Barcode sequencing to assess artemisinin response of strains. (A) Competition assay with continuous pressure of 10 nM DHA over 17 days. Supplementary Figure 2 shows a parallel experiment in which DHA was pulsed every 24 hours. (B) Experimental design for RSA-like experiment to assess artemisinin sensitivity. Barcoded strains were individually grown until schizont stages and combined at equal ratios followed by a density gradient (63% Percoll) enrichment of these stages. Next, 0-3 hour rings were synchronised with 5% Sorbitol and split in different replicates for the two conditions: Untreated (DMSO) and DHA (250nM). Genomic DNA was collected at 0, 3 and 72 hours and BarSeq libraries were generated as previously described. (C) RSA-like experiment showing growth of strains over the 72 hour assay. Y-axis shows the log2 of each barcode proportion normalised by the first time point (0 hours), followed by 3 and 72 hours (left panel) for three independent experiments. Right panel shows the change in proportion from 0 to 72 hours. Statistical significance between strains was calculated with one-way ANOVA and TukeyHSD.

We next adopted a modified version of the RSA using a lower DHA concentration (250 nM as opposed to 700 nM; Fig. 4B) to increase the relative survival of all strains and capture measurable barcodes, as the resistant strain forms only a small proportion of the total parasite pool. Using this experimental setup, BarSeq now revealed the expected response of the different strains to the drug, with CAM being the only line with increased survival, in concordance with the presence of the Kelch13-C580Y mutation and with those validated by other groups^2,19,29,30^ (Fig. 4C). Overall, this assay shows an approach for robust simultaneous phenotypic assessment of the drug response of multiple strains in a pool to the frontline antimalarial artemisinin, and offers the promise of ultimately screening hundreds of barcoded isolates on a single lane of Illumina MiSeq.

## Discussion

To accelerate efforts towards malaria elimination, novel antimalarials will be required to counter the current wave of multi-drug resistant parasites rapidly emerging. Having tools to better understand drug mode of action will be essential as we evaluate how new compounds function with current antimalarials, with a view to future combination therapies^31,32^. The complex interplay of drug resistance mutation acquisition and its overall impact on fitness has been highlighted in several studies^3,25,33–35^. However, being able to experimentally observe how drug-exposure interacts with particular genotypes to modulate overall parasite fitness has been challenging, and will better anticipate how these will perform in natural populations and polyclonal infections. In this work we present a method that could take us in that direction by tracking the competitive advantage of multiple *P. falciparum* strains, each carrying unique genomic barcodes, over time and drug exposure levels.

The use of BarSeq in *P. berghei* parasites using the *PlasmoGEM* vector resource has allowed functional profiling of gene knockouts at genome-scale^14^. As a complementary approach, we evaluated whether a similar approach could be applied to *P. falciparum* for understanding the fitness effects of drug resistance mutations *in vitro*, and their relevance in the current epidemiology of drug resistance. Unlike the more facile genetic manipulation of *P. berghei*, to do this in *P. falciparum* we used CRISPR-based editing to generate tagged lines that were uniquely barcoded at a constant safe-harbour genomic region, the pseudogene *rh3*. We transfected CRISPR-editing donors with 94 different barcodes within a single bulk transfection and subsequently obtained individually barcoded parasites via dilution cloning. Due to the low transfection efficiency of *P. falciparum* in general, and field isolates in particular, we only recovered a limited number of barcoded clones using this approach. Nonetheless, we generated sufficient clones to assay multiple strains in parallel, with replicates for two strains, and measure *in vitro* growth under a range of selection conditions. The sensitivity and throughput of the approach allowed us to observe that in the absence of drug pressure the more recent Cambodian isolate was unexpectedly equivalently fit to 3D7, which has been cultured for decades, whereas V1/S had a distinct fitness disadvantage to both strains. Application of increasing chloroquine pressure reversed the V1/S disadvantage, consistent with the presence of the mutant *pfcrt* allele, whereas mefloquine pressure restored the advantage of 3D7 over V1/S. Notably the CAM strain remained competitive under all conditions. These experiments confirm the impact of known drug resistance-associated mutations, as well as visualise inflection points where drug susceptibility and parasite fitness intersect.

Implementing this CRISPR-tagging and barcode sequencing approach to dissect fitness-conferring factors in the evolution to the frontline antimalarial artemisinin could have enormous value given the current wave of artemisinin resistant parasites that have emerged independently in different geographical locations^29,36,37^, and with questions remaining as to whether parasites in other regions of the world harbour a genetic background that might favour the emergence and spread of these resistance-associated mutations. By performing an RSA-like drug exposure for the pooled parasites in which early rings were pulsed with high concentration of DHA, we were able to detect enrichment of the Kelch13-mutant CAM parasite, but with the advantage that potentially tens or hundreds of strains could be profiled simultaneously unlike the standard RSA approach.

Several recent studies have developed genetic crosses of *P. falciparum* isolates using humanised mouse models^26^, aimed at understanding drug resistance determinants and how genetic background supports the resistance phenotype. Achieving this will require phenotyping of potentially hundreds of clonal progeny of resistant and sensitive parental strains, and changes in both drug sensitivity as well as in fitness measured by *in vitro* growth. One powerful method for identifying causal determinants in this context is bulk segregant analysis, which also operates using pooled parasites^26^. However, despite some up-front labour costs, barcode-tagging of parasites provides specific advantages in dissecting more complex polygenic traits by preserving haplotype information, while still allowing bulk analysis using assays such as competitive fitness and the artemisinin RSA to be tested simultaneously on large pools.

Studies in other systems such as the yeast model *Saccharomyces cerevisiae* have highlighted the potential resolution achievable by barcode sequencing. Levy et al 2015 demonstrated that up to 500,000 clonal lineages could be tracked over hundreds of generations, revealing the impact of accumulating mutations on shaping fitness landscapes^13^.

Recent years have seen an increase in the scale of adaptation of field isolates to culture^38,39^, and generating diversity via *in vitro* evolution of resistance to chemically diverse antimalarial compounds^40^. By barcoding and then assaying parasites encompassing a wide range of genomic diversity available in the field we should be able to understand how resistance mutations, compensatory changes, and genetic background interweave to shape the fitness landscape. Future studies will further expand the scope of tracking barcoded parasites to include other stages of the life cycle (e.g., transmission stages), as well as measuring growth under alternative host environments that will ultimately increase our understanding of biological fitness in *P. falciparum*.

## Methods

### Parasite cultures

Parasite strains used throughout this work were routinely cultured and propagated in human red blood cells provided by anonymous donors from the National Health Services Blood and Transplant (NHSBT). Informed consent was obtained by NHSBT as part of their recruitment process, and the use of RBCs was in accordance with relevant guidelines and regulations, with approval from the NHS Cambridgeshire Research Ethics Committee and the Wellcome Sanger Institute Human Materials and Data Management Committee. Culture media used throughout this work was as previously described^41^, with a gas mixture of 1% O_2_, 3% CO_2_ and 96% N_2_, and maintained at 37°C. Synchronisation of cultures was performed using 5%(w/v) sorbitol in water as previously described^42^ or with a Percoll gradient as previously described^43^. Laboratory-adapted parasite strains 3D7, V1/S were obtained from MR4, and clinical isolate PH0212-C (CAM) was isolated by Rick Fairhurst in Pursat, Cambodia^20^.

### Microscopy

Microscopy was performed with standard blood smears to determine parasitemia. A small aliquot of culture was smeared on a glass slide, fixed with 100% methanol, and stained with a 10% Giemsa solution (Sigma-Aldrich).

### Cloning of parasites by limiting dilution

Parasites recovered after transfection were cloned using a limiting dilution protocol modified from^44^. Parasites were diluted into a 96-well plate at 0.5-0.8 parasites per well at 1.8% haematocrit, for a recovery of 50% of the plate corresponding to clonal parasites. For rapid screening of positive wells, a DNA stain SYBR Green method was used, and both empty wells or those with much higher fluorescence than the average (more than one clone at start) were discarded.

### Barcode sequencing

The barcoding approach was adapted for *P. falciparum* from PlasmoGEM^45^. We recently described successful transfection of episomally-maintained pools of barcoded vectors^17^. Here we used these pools together with CRISPR/Cas9-expressing vectors for barcode integration into *Rh3* (PF3D7_1252400). In brief, the barcode-containing region from 96 different pGEM bacterial stocks was amplified simultaneously after individually growing and pooling them (TB medium with 30 μg/mL of kanamycin). Primers p212 and p219 (Supplementary Table 4) were designed to amplify the 120 bp barcode amplicon with overlap sequences to a Nhel/Ncol digested *P. falciparum* vector pCC1^16^. Gibson cloning ^46^ was used for integration of amplicon regions in-between two homology regions of *Pfrh3* (Fig. 1A), each spanning approximately 1kb.

CRISPR/Cas9 plasmids were generated using a pDC2 vector backbone as previously described^47^, expressing Cas9 under a *P. falciparum-specific* promoter and a single guide RNA under a U6 promoter for RNA Pol III^19^. We replaced the hDHFR positive selection cassette for yDHOH ^48^ in order to select for both co-transfected plasmids.

### Parasite transfections

Transfections were performed using the pool of 94 barcoded vectors as well as the CRISPR/Cas9-expressing vectors on ring-stages at 5-10% parasitemia. A Lonza Nucleofactor 4D was used with the programme P3-CM150. Plasmid DNA composed of a 1:1 mixture of the targeting and the donor vectors was resuspended in 100 μL of P3 buffer (Lonza), containing 3 μL ATP (625 mM). Parasites were centrifuged and the DNA solution was used to resuspend 100 μL of packed RBCs. Selection with WR99210 was applied 24 hours post-transfection, with 5 nM being applied for 3D7, and 10nM for V1/S and CAM. Parasites recovered after transfection were cloned by limiting dilution^44^, and genotyped prior to confirming barcode integration through sequencing.

### Barcode sequencing

Uniquely barcoded clones were expanded and independently synchronised by a Percoll gradient to enrich for late segmenting schizonts, followed by sorbitol synchronisation^42^. Parasitemia was adjusted by using flow cytometry and mixtures were made at equal starting parasitemia on day zero. Cultures were diluted to maintain parasitemia between 1-5% throughout the time course, which would define the day of collection. Assays were conducted in duplicate or triplicate, each on 10 mL culture volumes, and at different timepoints cells were collected and lysed with 0.05% saponin for genomic DNA (gDNA) isolation. Genomic DNA was extracted using a Qiagen Blood and Tissue kit. 50ng of extracted gDNA was used as a template for a nested PCR reaction using KAPA HiFi 2X polymerase master mix. A first PCR reaction was performed to amplify from outside the homology regions of *Pfrh3* in order to avoid amplification from stably maintained episomes, using primers p191 and p194, followed by a nested 2-step PCR reaction using p1356 and p1357 containing adapters for Illumina sequencing (PCR1). See Supplementary Table 4 for primer sequences. Following adapter ligation, 5 μL of PCR1 were used with paired-end index primers (Nextera XT) for PCR2. PCR products were purified from this last reaction only, using a Macherey Nagel PCR purification kit and eluted DNA was quantified using Qubit broad sensitivity kit, multiplexed and diluted to a final concentration of 4 nM. Samples were loaded onto an Illumina MiSeq sequencer, using a MiSeq Reagent Kit v2 (300 cycle). They were loaded at a low cluster density (<400k cluster density), and 50% of PhiX was spiked in, as described in Gomes et al. (2015) for low complexity libraries ^45^.

### Barcode counting and analysis

Barcode counting was performed as in (Gomes et al 2015). The output Illumina fastq reads from the MiSeq, represented by unique index tags corresponding to the different time points, were separated and analysed with a script that identified correct flanking sequences and counted exact matches of unique barcodes between these regions.

## Data availability

All data necessary to perform the analysis of the barcode sequencing of parasite strains is available as part of the supplementary material (Supplementary Tables 2 and 3).

## Acknowledgements

We would like to thank current and former members of the Lee, Rayner, and Billker labs for constructive feedback. We are grateful to Thanat Chookajorn and Charin Modchang from Mahidol University (Bangkok, Thailand) for productive discussions on the project. We are also grateful to the staff in Sanger Scientific Operations for their support with sequencing. This work was supported by funding from Wellcome (206194).

## Author contributions

M.C., J.R. and M.L. conceived the study. M.C. performed the barcode cloning and transfection for *P. falciparum*, M.C., N.D., and M.R. performed the barseq assays, M.C. and T.S. analysed the data, N.D., M.R. and M.L. performed the drug assays, Z.Z. constructed the Cas9-yDHODH vector. ML and JR supervised the study. All authors contributed to writing the paper.

## Competing interests

The authors declare no competing interests.

---

**Supplementary Figure 1.**
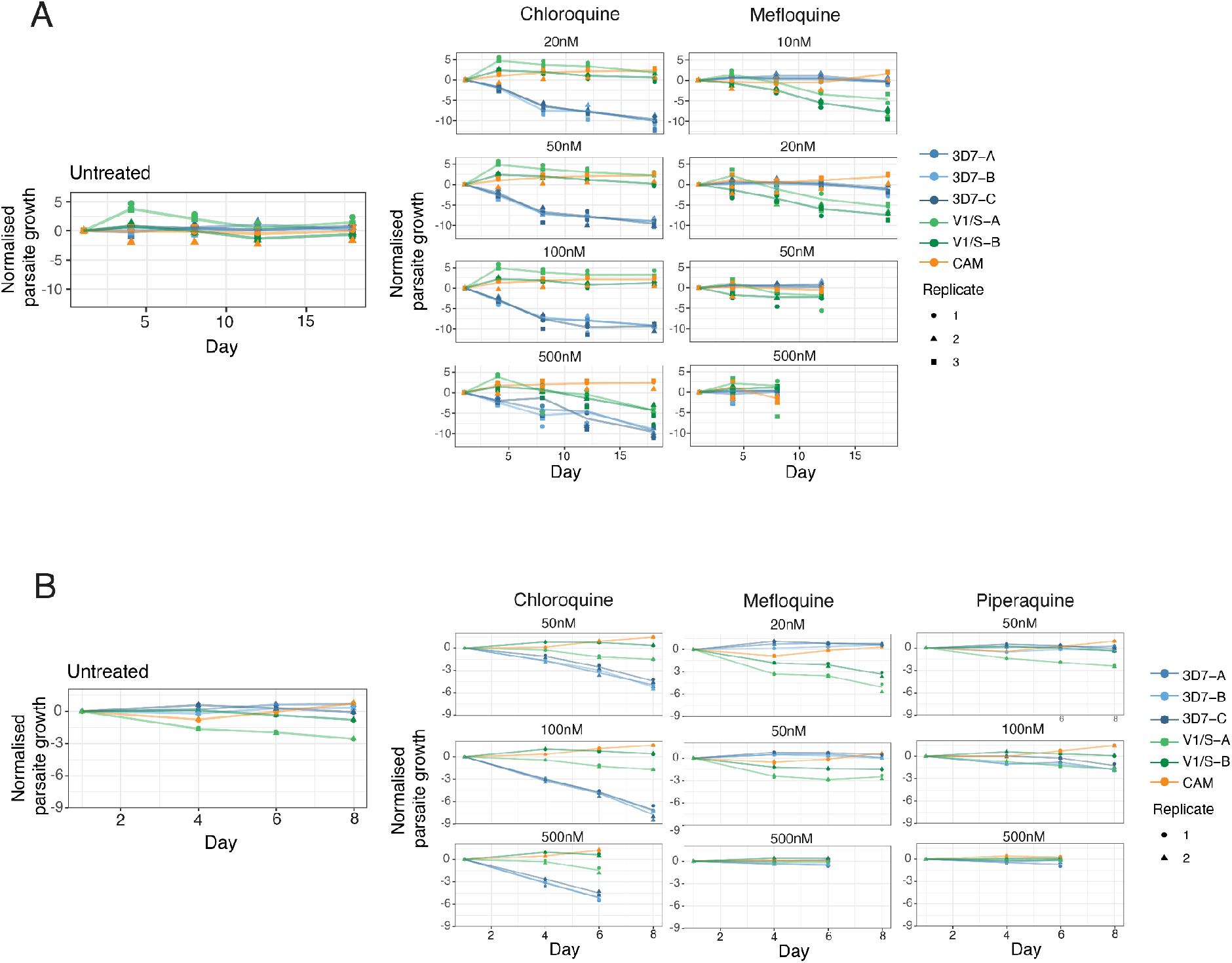
Drug response of barcoded strains at different concentrations in two independent experiments. (A) Normalised growth (y-axis) in the untreated control and at increasing concentrations of chloroquine (left) starting at the IC_50_ for 3D7 (20nM), 50, 100 and 500nM over 17 days. Right panel shows normalised growth in presence of mefloquine at concentrations ranging from 10nM (IC_50_ for V1/S, the most sensitive strain), 20 and 50nM over 17 days and a time course lasting 8 days at the highest concentration (500nM). (B) Normalised growth (y-axis) in the untreated control and increasing concentrations of chloroquine, mefloquine and piperaquine. For chloroquine (left), 50 and 100nM were used over a period of 8 days, for mefloquine (middle) 20 and 50nM over 8 days and piperaquine 50 and 100nM for 8 days (right). 500nM treatments for all three antimalarial drugs were performed for 6 days.

**Supplementary Figure 2.**
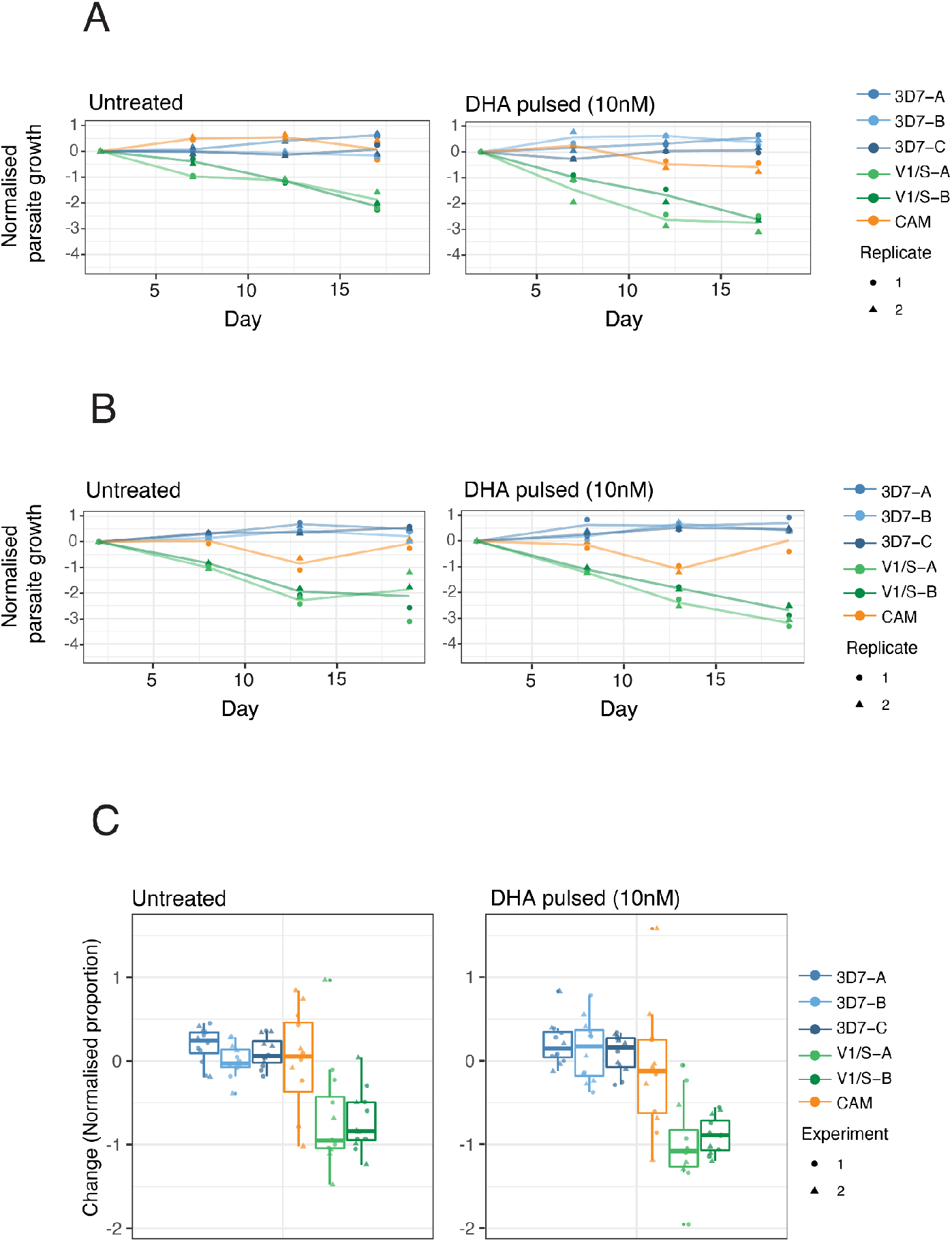
Competition assay using DHA, pulsing every 24 hours for 3 hours at 10 nM, for two independent experiments. To complement Figure 4. (A-B) Normalised growth (y-axis) in the untreated control and at 10 nM DHA for the barcoded pool of strains for two independent experiments (four biological replicates). (C) Log2-change in normalised growth (y-axis) for the untreated control of four biological replicates shown in A and B.

**Supplementary Table 1:** IC_50_ data for 3D7, V1/S and CAM.

| **Antimalarial** | **3D7** |  |  | **V1/S** |  |  | **CAM** |  |  |
| --- | --- | --- | --- | --- | --- | --- | --- | --- | --- |
|  | IC50 (nM) | Std Dev | N | IC50 (nM) | Std Dev | N | IC50 (nM) | Std Dev | N |
| Chloroquine | 23.2 | 6.8 | 3 | 332.4 | 57.2 | 3 | 576.4 | 64.3 | 3 |
| Mefloquine | 32.7 | 4.8 | 3 | 7.2 | 3.8 | 3 | 29.2 | 5.2 | 3 |
| Piperaquine | 5.4 | 0.9 | 6 | 10.2 | 1.2 | 6 | 10.4 | 1.9 | 6 |
| Dihydroartemisinin | 2.0 | 0.2 | 3 | 1.6 | 0.6 | 3 | 3.1 | 1.4 | 3 |

## References

1. Straimer, J. et al. K13-propeller mutations confer artemisinin resistance in *Plasmodium falciparum* clinical isolates. Science 347, (2015).

2. Straimer, J. et al. Plasmodium falciparum K13 Mutations Differentially Impact Ozonide Susceptibility and Parasite Fitness In Vitro. MBio 8, (2017).

3. Nair, S. et al. Do fitness costs explain the rapid spread of kelch13-C580Y substitutions conferring artemisinin resistance? Antimicrob. Agents Chemother. 62, (2018)

4. Amato, R. et al. Origins of the current outbreak of multidrug-resistant malaria in southeast Asia: a retrospective genetic study. Lancet Infect. Dis. 18, 337–345 (2018).

5. Payne, D. Spread of chloroquine resistance in *Plasmodium falciparum*. Parasitol. Today 3, 241–246 (1987).

6. Griffing, S. et al. pfmdr1 amplification and fixation of pfcrt chloroquine resistance alleles in Plasmodium falciparum in Venezuela. Antimicrob. Agents Chemother. 54, 1572–1579 (2010).

7. Mita, T., Tanabe, K. & Kita, K. Spread and evolution of *Plasmodium falciparum* drug resistance. Parasitol. Int. 58, 201–209 (2009).

8. Pearce, R. J. et al. Multiple origins and regional dispersal of resistant dhps in African Plasmodium falciparum malaria. PLoS Med. 6, e1000055 (2009).

9. Kublin, J. G. et al. Reemergence of chloroquine-sensitive *Plasmodium falciparum* malaria after cessation of chloroquine use in Malawi. J. Infect. Dis. 187, 1870–1875 (2003).

10. Petersen, I. et al. Balancing drug resistance and growth rates via compensatory mutations in the *Plasmodium falciparum* chloroquine resistance transporter. Mol. Microbiol. 97, 381–395 (2015).

11. Baragaña, B. et al. A novel multiple-stage antimalarial agent that inhibits protein synthesis. Nature 522, 315–320 (2015).

12. Hayward, R., Saliba, K. J. & Kirk, K. pfmdr1 mutations associated with chloroquine resistance incur a fitness cost in *Plasmodium falciparum*. Mol. Microbiol. 55, 1285–1295 (2005).

13. Levy, S. F. et al. Quantitative evolutionary dynamics using high-resolution lineage tracking. Nature 519, 181–186 (2015).

14. Bushell, E. et al. Functional Profiling of a *Plasmodium* Genome Reveals an Abundance of Essential Genes. Cell 170, 260–272.e8 (2017).

15. Sidik, S. M. et al. A Genome-wide CRISPR Screen in Toxoplasma Identifies Essential Apicomplexan Genes. Cell 166, 1423–1435.e12 (2016).

16. Maier, A. G. et al. Exported proteins required for virulence and rigidity of *Plasmodium falciparum-infected human erythrocytes*. Cell 134, 48–61 (2008).

17. Carrasquilla, M. et al. Defining multiplicity of vector uptake in transfected *Plasmodium* parasites. Sci. Rep. 10, 10894 (2020).

18. Lim, M. Y.-X. et al. UDP-galactose and acetyl-CoA transporters as Plasmodium multidrug resistance genes. Nat Microbiol 1, 16166 (2016).

19. Ghorbal, M. et al. Genome editing in the human malaria parasite *Plasmodium falciparum* using the CRISPR-Cas9 system. Nat. Biotechnol. 32, 819–821 (2014).

20. Lee, A. H. & Fidock, D. A. Evidence of a Mild Mutator Phenotype in Cambodian *Plasmodium falciparum* Malaria Parasites. PLoS One 11, e0154166 (2016).

21. Vijaykadga, S. et al. In vivo sensitivity monitoring of mefloquine monotherapy and artesunate--mefloquine combinations for the treatment of uncomplicated falciparum malaria in Thailand in 2003. Trop. Med. Int. Health 11, 211–219 (2006).

22. Denis, M. B. et al. Surveillance of the efficacy of artesunate and mefloquine combination for the treatment of uncomplicated falciparum malaria in Cambodia. Trop. Med. Int. Health 11, 1360–1366 (2006).

23. van der Pluijm, R. W. et al. Determinants of dihydroartemisinin-piperaquine treatment failure in *Plasmodium falciparum* malaria in Cambodia, Thailand, and Vietnam: a prospective clinical, pharmacological, and genetic study. Lancet Infect. Dis. (2019).

24. Ménard, D. & Fidock, D. A. Accelerated evolution and spread of multidrug-resistant *Plasmodium falciparum* takes down the latest first-line antimalarial drug in southeast Asia. The Lancet Infectious Diseases (2019) doi:10.1016/s1473-3099(19)30394-9.

25. Li, X. et al. Genetic mapping of fitness determinants across the malaria parasite *Plasmodium falciparum* life cycle. PLoS Genet. 15, e1008453 (2019).

26. Vendrely, K. M., Kumar, S., Li, X. & Vaughan, A. M. Humanized Mice and the Rebirth of Malaria Genetic Crosses. Trends Parasitol. 36, 850–863 (2020).

27. Witkowski, B. et al. Novel phenotypic assays for the detection of artemisinin-resistant *Plasmodium falciparum* malaria in Cambodia: in-vitro and ex-vivo drug-response studies. Lancet Infect. Dis. 13, 1043–1049 (2013).

28. Noedl, H. et al. Evidence of Artemisinin-Resistant Malaria in Western Cambodia. N. Engl. J. Med. 359, 2619–2620 (2008).

29. Ariey, F. et al. A molecular marker of artemisinin-resistant *Plasmodium falciparum* malaria. Nature 505, (2014).

30. Phyo, A. P. et al. Poor response to artesunate treatment in two patients with severe malaria on the Thai–Myanmar border. Malar. J. 17, 30 (2018).

31. Burrows, J. N. et al. New developments in anti-malarial target candidate and product profiles. Malar. J. 16, 26 (2017).

32. Nations, United. The Millenium Developmental Goals Report. United Nations (2015).

33. Stokes, B. H. et al. Plasmodium falciparum K13 mutations in Africa and Asia impact artemisinin resistance and parasite fitness. Elife 10, (2021).

34. Bunditvorapoom, D. et al. Fitness Loss under Amino Acid Starvation in Artemisinin-Resistant *Plasmodium falciparum* Isolates from Cambodia. Sci. Rep. 8, 12622 (2018).

35. Tirrell, A. R. et al. Pairwise growth competitions identify relative fitness relationships among artemisinin resistant *Plasmodium falciparum* field isolates. Malar. J. 18, 295 (2019).

36. Miotto, O. et al. Emergence of artemisinin-resistant *Plasmodium falciparum* with kelch13 C580Y mutations on the island of New Guinea. PLoS Pathog. 16, e1009133 (2020).

37. Mathieu, L. C. et al. Local emergence in Amazonia of *Plasmodium falciparum* k13 C580Y mutants associated with in vitro artemisinin resistance. Elife 9, (2020).

38. Claessens, A., Affara, M., Assefa, S. A., Kwiatkowski, D. P. & Conway, D. J. Culture adaptation of malaria parasites selects for convergent loss-of-function mutants. Sci. Rep. 7, 41303 (2017).

39. Nsobya, S. L., Kiggundu, M., Joloba, M., Dorsey, G. & Rosenthal, P. J. Complexity of *Plasmodium falciparum* clinical samples from Uganda during short-term culture. J. Infect. Dis. 198, 1554–1557 (2008).

40. Cowell, A. N. et al. Mapping the malaria parasite druggable genome by using in vitro evolution and chemogenomics. Science 359, 191–199 (2018).

41. Trager, W. & Jensen, J. B. Human malaria parasites in continuous culture. Science 193, 673–675 (1976).

42. Radfar, A. et al. Synchronous culture of *Plasmodium falciparum* at high parasitemia levels. Nat. Protoc. 4, 1899–1915 (2009).

43. Knuepfer, E., Napiorkowska, M., van Ooij, C. & Holder, A. A. Generating conditional gene knockouts in *Plasmodium* – a toolkit to produce stable DiCre recombinase-expressing parasite lines using CRISPR/Cas9. Scientific Reports 7, (2017).

44. Rosario, V. Cloning of naturally occurring mixed infections of malaria parasites. Science 212, 1037–1038 (1981).

45. Gomes, A. R. et al. A genome-scale vector resource enables high-throughput reverse genetic screening in a malaria parasite. Cell Host Microbe 17, 404–413 (2015).

46. Gibson, D. G. et al. Complete chemical synthesis, assembly, and cloning of a *Mycoplasma genitalium* genome. Science 319, 1215–1220 (2008).

47. Adjalley S, L. M. CRISPR/Cas9 editing of the *Plasmodium falciparum* genome. in Methods in Molecular Biology (2022).

48. Ganesan, S. M. et al. Yeast dihydroorotate dehydrogenase as a new selectable marker for *Plasmodium falciparum* transfection. Mol. Biochem. Parasitol. 177, 29–34 (2011).

